# Chlamydial Histones Control Developmental Fitness in the Next Infection Cycle

**DOI:** 10.64898/2026.02.10.705219

**Authors:** Yuxuan Wang, Matthew Pan, Temitope V. Coker, Jing Wang, Lingling Wang, Guangming Zhong, Huizhou Fan

**Author notes:** Correspondence: Huizhou Fan, PhD.

## Abstract

The unique chlamydial developmental cycle comprises three stages: primary differentiation of infectious elementary bodies (EBs) into reticulate bodies (RBs), RB replication, and secondary differentiation into progeny EBs. Extensive chromosome remodeling during RB-to-EB differentiation is thought to be mediated by the histones HctA and HctB. Here, we used an inducible CRISPR interference system to repress *hctA, hctB*, or both genes during development in *Chlamydia trachomatis*. Surprisingly, repression of either histone gene alone or in combination caused only modest reductions in EB yield and did not prevent nucleoid condensation during the parental developmental cycle. In contrast, when progeny EBs generated under histone-repressing conditions were used to initiate secondary infections in the absence of inducer, histone deficiency during EB maturation profoundly impaired fitness in the next infection cycle. Secondary cultures initiated with HctA-deficient EBs exhibited a delayed onset of genome replication, consistent with inefficient primary EB-to-RB differentiation, whereas combined repression of *hctA* and *hctB* caused both delayed genome replication and persistently reduced genome accumulation, indicative of defects in RB formation and subsequent growth. Repression of *hctB* alone did not measurably affect genome replication in secondary cultures. Together, these findings reveal a transgenerational role for chlamydial histones and establish chromosome organization during EB maturation as a key determinant of developmental fitness across infection cycles.

**IMPORTANCE:** Chlamydial histones HctA and HctB are unusual among bacterial chromatin-binding proteins in that they share sequence homology with mammalian histones and are developmentally regulated during the formation of infectious particles. Here, we show that reduced expression of HctA and HctB has only modest effects on genome condensation and EB production, consistent with partial functional redundancy between the two histones and suggesting that additional chromatin factors contribute to EB chromosome compaction. In contrast, deficiency of HctA and HctB during EB maturation has profound consequences in the next infection cycle, impairing primary EB-to-RB differentiation and subsequent RB growth. These findings reveal a previously unrecognized transgenerational role for chlamydial histones and establish chromosome organization during EB maturation as a key determinant of developmental fitness across infection cycles.

## INTRODUCTION

*Chlamydia trachomatis* is an obligate intracellular bacterium whose pathogenic success depends on a biphasic developmental cycle that alternates between two morphologically and physiologically distinct forms: the infectious elementary body (EB) and the replicative reticulate body (RB) [1-3]. EBs are small, electron-dense particles with highly compacted nucleoids, adapted for extracellular survival and host-cell entry, whereas RBs are larger and fragile, with diffuse chromatin consistent with active transcription and DNA replication [1-3]. Following internalization, EBs differentiate into RBs within a membrane-bound inclusion, undergo multiple rounds of division, and subsequently redifferentiate into progeny EBs that are released to infect new host cells [1-6].

The dramatic chromatin remodeling that accompanies RB-to-EB differentiation is thought to be mediated by two developmentally regulated histones, HctA (Hc1) and HctB (Hc2) [7-11]. Both proteins bind DNA and repress transcription *in vitro* [12-14], and their abundance increases late in the developmental cycle as EBs form [15, 16]. However, the two genes are regulated differently. *hctA* transcripts accumulate earlier in infection but are translationally silenced by the small RNA *ihtA* until late stages, whereas *hctB* is expressed primarily during terminal RB-to-EB differentiation [17, 18].

Early heterologous-expression studies revealed strikingly distinct phenotypes for the two proteins. Overproduction of HctA in *Escherichia coli* causes rapid nucleoid condensation and severe growth inhibition, whereas HctB overexpression induces the formation of unusual coil-like DNA structures whose relevance to chlamydial chromosome organization remains unclear [7, 8, 10]. These divergent phenotypes, together with the distinct developmental regulation of the two genes, suggested that HctA and HctB might perform non-identical biological functions in *Chlamydia* itself, functions that could not be resolved until genetic tools became available to perturb histone expression directly in the native pathogen.

Here, we use inducible CRISPRi to knock down *hctA, hctB*, or both genes to investigate whether and how chlamydial histones regulate EB formation and developmental fitness. Surprisingly, repression of either histone gene alone or in combination has only limited effects on EB morphogenesis in the parental cycle, whereas deficiency of HctA alone, or in combination with HctB, compromises fitness in the subsequent infection cycle, with the dual knockdown producing the most severe phenotype. These results indicate that HctA and HctB perform overlapping yet distinct functions during chlamydial development and reveal an unexpected transgenerational role for histone-mediated chromosome organization. They further suggest that chromatin architecture established during EB maturation influences the efficiency of developmental reactivation following reinfection.

## RESULTS

### CRISPRi enables efficient and specific knockdown of *hctA* or *hctB*

To examine the roles of chlamydial histones during late development, we generated *C. trachomatis* L2 strains L2/hctA-iKD and L2/hctB-iKD, which express ATC-inducible ddCas12 and guide RNAs targeting either *hctA* or *hctB*. To determine knockdown efficiency, anhydrotetracycline (ATC) was added at 0 hour postinoculation (hpi), and transcript levels were quantified at 26 hpi, corresponding to the early late developmental phase. Under these conditions, ATC induction increased ddCas12 transcript abundance by approximately six-fold, resulting in strong knockdown of the targeted histone genes: *hctA* and *hctB* transcripts were reduced by 94% and 84% in L2/hctA-iKD and L2/hctB-iKD cultures, respectively, relative to ATC-free controls (Fig. 1A). Silencing of one histone gene did not affect expression of the other and did not alter levels of *ihtA*, a small RNA that binds hctA mRNA to inhibit its translation (Fig. 1B). In the non-targeting control L2/ddCas12-ntg strain, ATC induced ddCas12 expression without affecting histone transcript abundance (Fig. 1C). Collectively, these results demonstrate that CRISPRi enables selective and robust knockdown of individual histone genes during late development.

**Figure 1.**
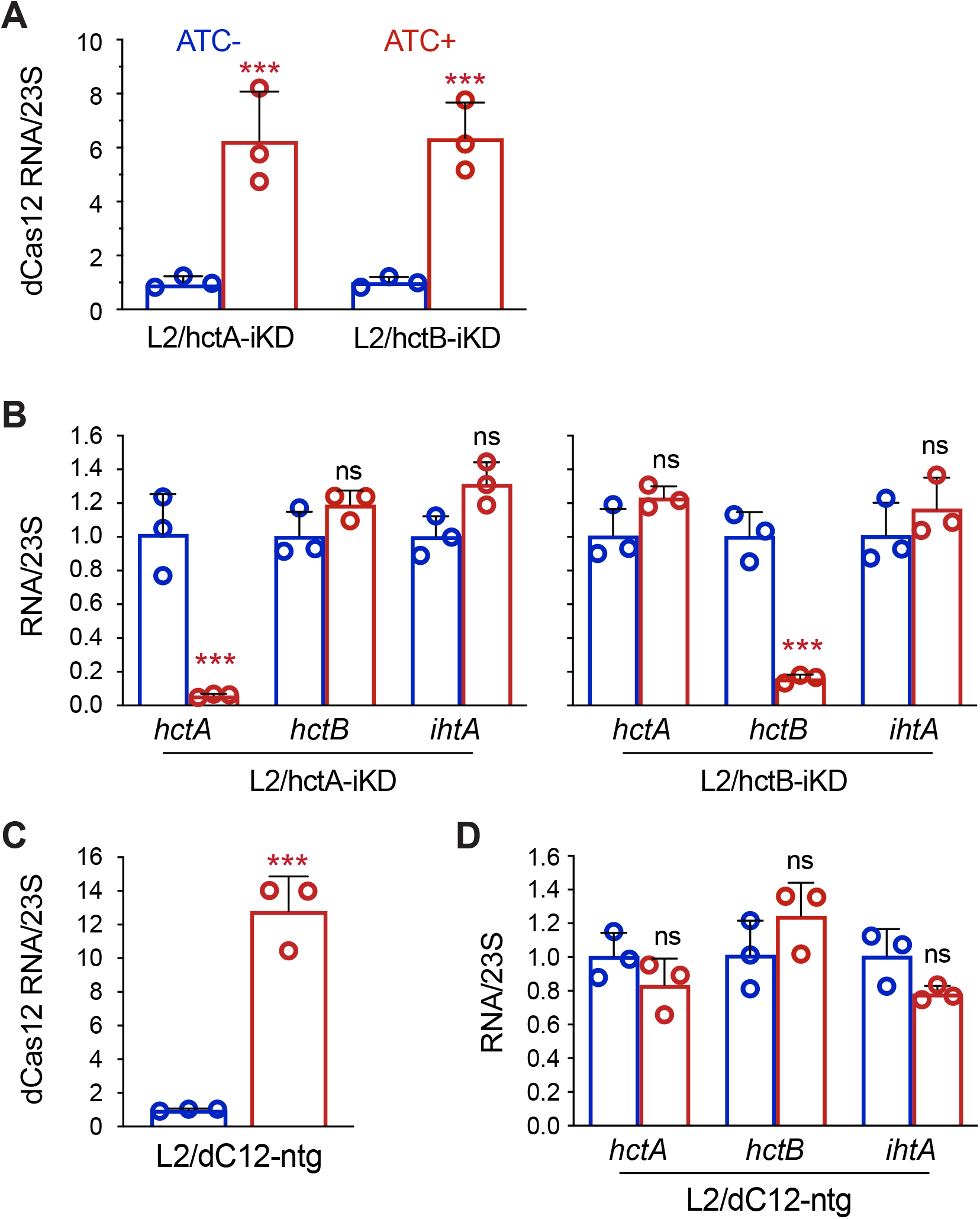
ATC induces ddCas12 expression and selectively reduces histone transcripts in L2/hctA-iKD and L2/hctB-iKD. (A) ATC-induced ddCas12 mRNA levels in L2/hctA-iKD and L2/hctB-iKD. (B) ATC-dependent reduction of the targeted histone transcript in L2/hctA-iKD and L2/hctB-iKD, with no effect on the non-targeted histone or *ihtA*. (C) ATC-induced ddCas12 mRNA levels and unchanged *hctA, hctB*, and *ihtA* transcripts in the control strain L2/ddCas12-ntg. (A-C) ATC was added at 0 hpi, total RNA was isolated at 24 hpi, and transcript abundance was determined by RT-qPCR and normalized to 23S rRNA. Data represent averages ± standard deviations from three independent biological replicates, with individual points shown. Statistical significance relative to ATC-free cultures is indicated (***, *P* < 0.001; ns, not significant).

### Single-gene histone knockdowns do not impair RB replication but modestly reduce EB production

Because HctA and HctB are developmentally regulated and implicated primarily in late-stage genome condensation [7, 8], we predicted that their knockdown would not strongly affect RB replication but would compromise EB production. Indeed, in cultures grown with ATC, the genome replication kinetics of both L2/hctA-iKD and L2/hctB-iKD were indistinguishable from those of ATC-free controls (Fig. 2). Surprisingly, IFU assays revealed that ATC-induced knockdown in L2/hctA-iKD and L2/hctB-iKD had modestly reduced EB yields, typically by only two-to three-fold (Fig. 3). These effects were far weaker than those reported for inhibition of essential late developmental regulators [19, 20], suggesting that *Chlamydia* tolerates large reductions in histone expression during EB biogenesis.

**Fig. 2.**
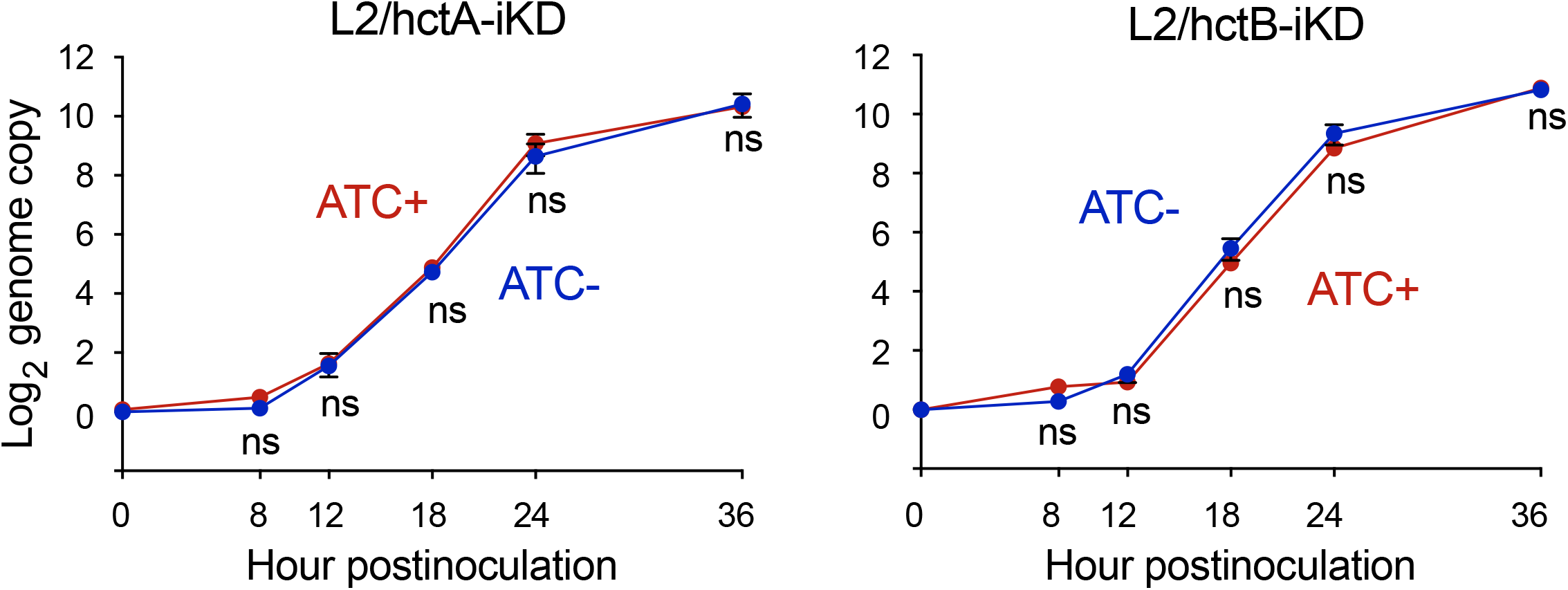
Genome replication is unaffected by ATC-induced *hctA* or *hctB* knockdown. Genome copy number was determined by qPCR at the indicated time points in L2/hctA-iKD and L2/hctB-iKD cultures with or without ATC. Data represent averages ± standard deviations from three independent biological replicates. Differences between ATC-treated and ATC-free cultures were not statistically significant at any time point for either strain.

**Fig. 3.**
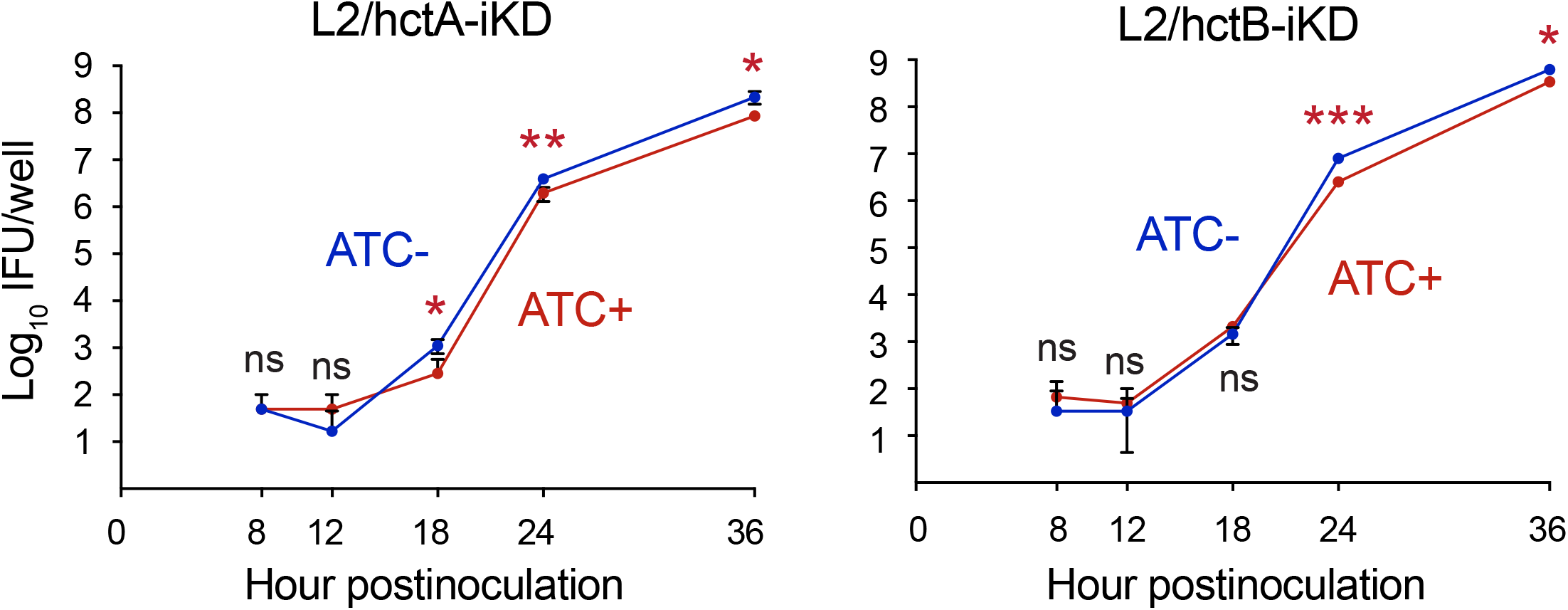
EB production is modestly reduced by ATC-induced *hctA* or *hctB* knockdown. Infectious progeny were measured by IFU assays at the indicated time points in L2/hctA-iKD and L2/hctB-iKD cultures with or without ATC. Data represent averages ± standard deviations from three independent biological replicates. Levels of statistical significance for differences between ATC-treated and ATC-free cultures are indicated (*, **, and *** signify *P* < 0.05, 0.01, and 0.001, respectively).

### Histone gene knockdowns do not prevent EB morphogenesis or nucleoid condensation

Because IFU assays indicated that histone knockdown still permitted production of infectious progeny, we asked whether these particles corresponded to morphologically authentic EBs with condensed genomes. ATC-free and ATC-treated cultures of L2/hctA-iKD and L2/hctB-iKD were therefore examined by transmission electron microscopy at 36 hpi. Similar to ATC-free cultures, ATC-treated cultures contained both RBs, characterized by larger size and relatively low electron density, and EBs with smaller size and high electron density (Fig. 4). These findings indicate that knockdown of either histone gene alone does not abolish genome condensation or EB morphogenesis during late developmental stages, consistent with the modest reductions in EB yield measured by IFU assays.

**Fig. 4.**
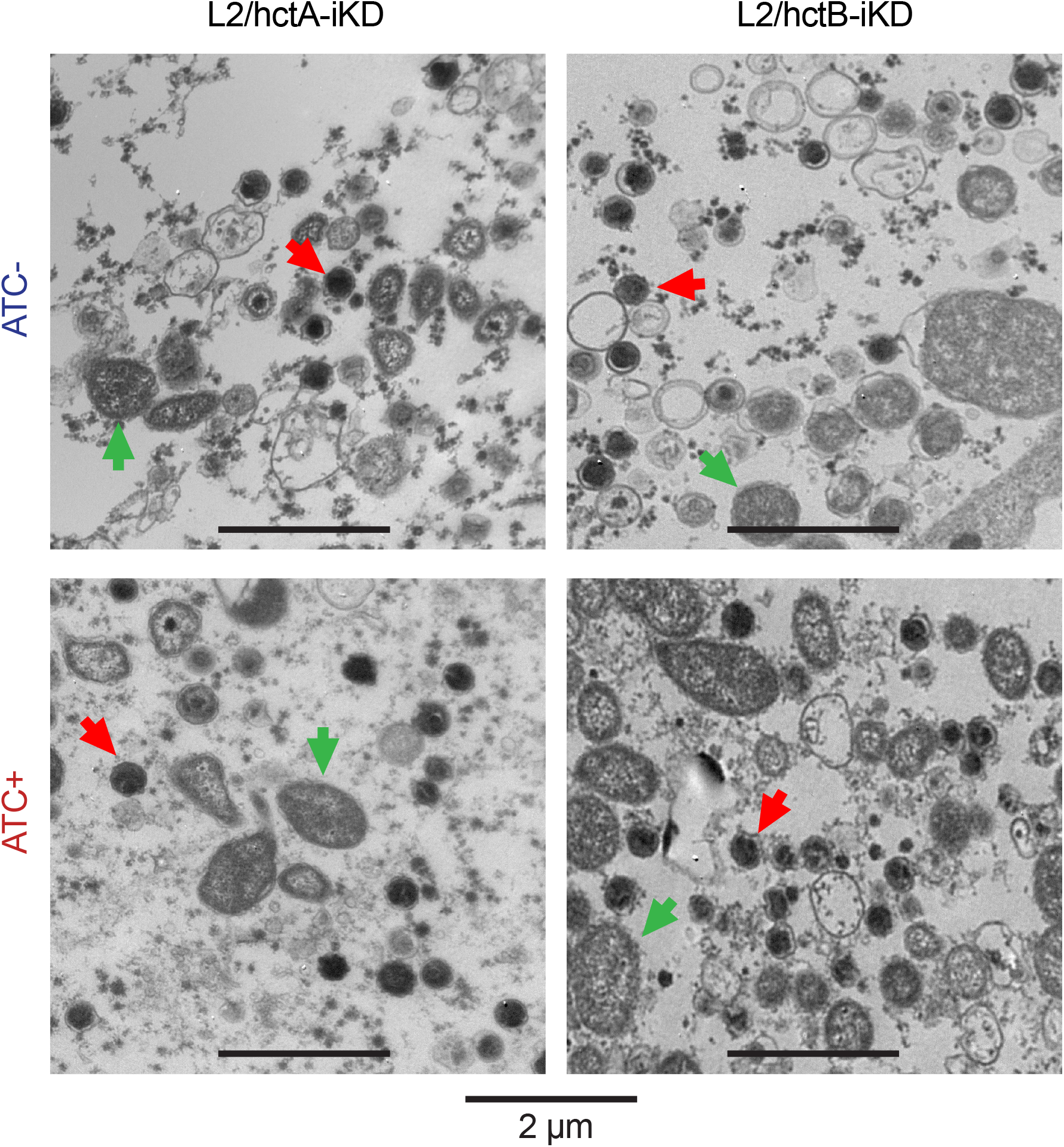
EB nucleoid condensation is preserved after ATC-induced hctA or hctB knockdown. Representative transmission electron micrographs of L2/hctA-iKD and L2/hctB-iKD cultures grown with or without ATC. ATC was added at 0 hpi, and samples were collected at 36 hpi for TEM analysis. Representative RBs and EBs are indicated in the images.

### Knockdown of both hctA and hctB still has a limited effect on EB formation

The limited phenotypes observed after single-gene knockdown (Figs. 2–4) suggested that HctA and HctB might perform overlapping functions during EB formation. To test redundancy, we generated a dual-targeting strain (L2/hctAB-iKD) expressing guide RNAs targeting both histone genes. ATC treatment reduced hctA and hctB transcript levels by 97% and 88%, respectively (Fig. 5A).

**Fig. 5.**
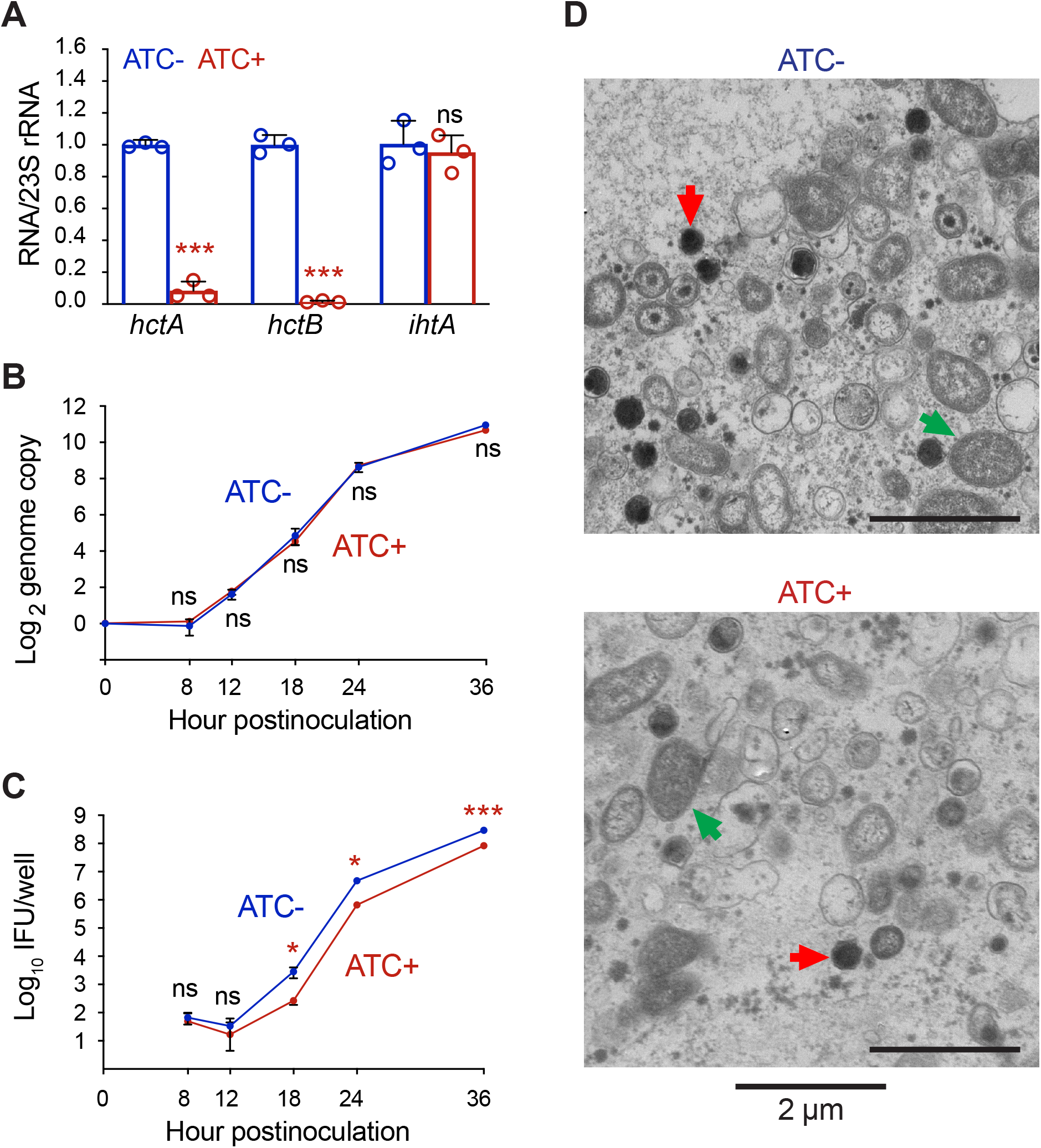
Fig. 5. Knockdown of both *hctA* and *hctB* causes limited EB production reduction while preserving nucleoid condensation. (A) ATC-dependent reduction of *hctA* and *hctB* transcripts but not *ihtA* levels in L2/hctAB-iKD. (B) Genome copy number over time in L2/hctAB-iKD cultures grown with or without ATC. (C) EB production measured by IFU assays in L2/hctAB-iKD cultures with or without ATC. (D) Representative transmission electron micrographs of progeny EBs from ATC-free and ATC-treated L2/hctAB-iKD cultures at 36 hpi, with RBs and EBs indicated. (A-C) ATC was added at 0 hpi. RNA for RT-qPCR was isolated at 24 hpi. Data represent averages ± standard deviations from three independent biological replicates. Statistical significance between ATC-treated and ATC-free cultures is indicated in the graphs.

Similar to the single-gene knockdowns in L2/hctA-iKD and L2/hctB-iKD, genome replication kinetics in L2/hctAB-iKD remained indistinguishable between ATC-free and ATC-treated cultures (Fig. 5B). EB yields in ATC-treated L2/hctAB-iKD cultures were reduced slightly more than in ATC-treated L2/hctA-iKD and L2/hctB-iKD; however, the decreases remained less than ten-fold at each time point examined (Fig. 5C), supporting the conclusion that the effect of combined histone gene knockdown on EB formation was still modest. Transmission electron microscopy further showed that progeny EBs in L2/hctAB-iKD cultures retained condensed nucleoids under dual-knockdown conditions (Fig. 5D).

Together, these data indicate that simultaneous knockdown of both histone genes does not substantially exacerbate the phenotypes observed after single-gene knockdown and that RB replication and EB morphogenesis remain largely intact despite severe reductions in histone transcript levels.

### Histone deficiency during EB formation compromises development in the next cycle

Although histone knockdown caused only limited defects during the initial developmental cycle, we noticed that inclusions were consistently smaller during IFU assays when secondary cultures were inoculated with EBs harvested from ATC+ L2/hctA-iKD and ATC+ L2/hctAB-iKD primary infections, even though all secondary cultures were grown without ATC. This observation prompted us to examine systematically whether histone deficiency during EB formation impaired the ability of progeny EBs from all three knockdown strains to initiate subsequent infections by analyzing inclusion area, mKate fluorescence, and genome replication kinetics in secondary cultures.

Representative fluorescence images at 18 and 36 h postinoculation showed that inclusions formed by ATC+ L2/hctA-iKD EBs were smaller and dimmer than those formed by ATC− L2/hctA-iKD EBs, whereas inclusions formed by ATC+ L2/hctB-iKD EBs appeared largely similar to the corresponding ATC− controls (Fig. 6A). Inclusions formed by ATC+ L2/hctAB-iKD EBs were markedly smaller and dimmer than ATC− controls. Quantification confirmed that the reductions in inclusion area and mKate intensity were more severe for cultures inoculated with ATC+ L2/hctAB-iKD EBs than for those inoculated with ATC+ L2/hctA-iKD EBs, while ATC+ L2/hctB-iKD EBs produced only a modest, transient decrease in inclusion area at 18 h with no detectable reduction in mKate intensity (Fig. 6B).

**Fig. 6.**
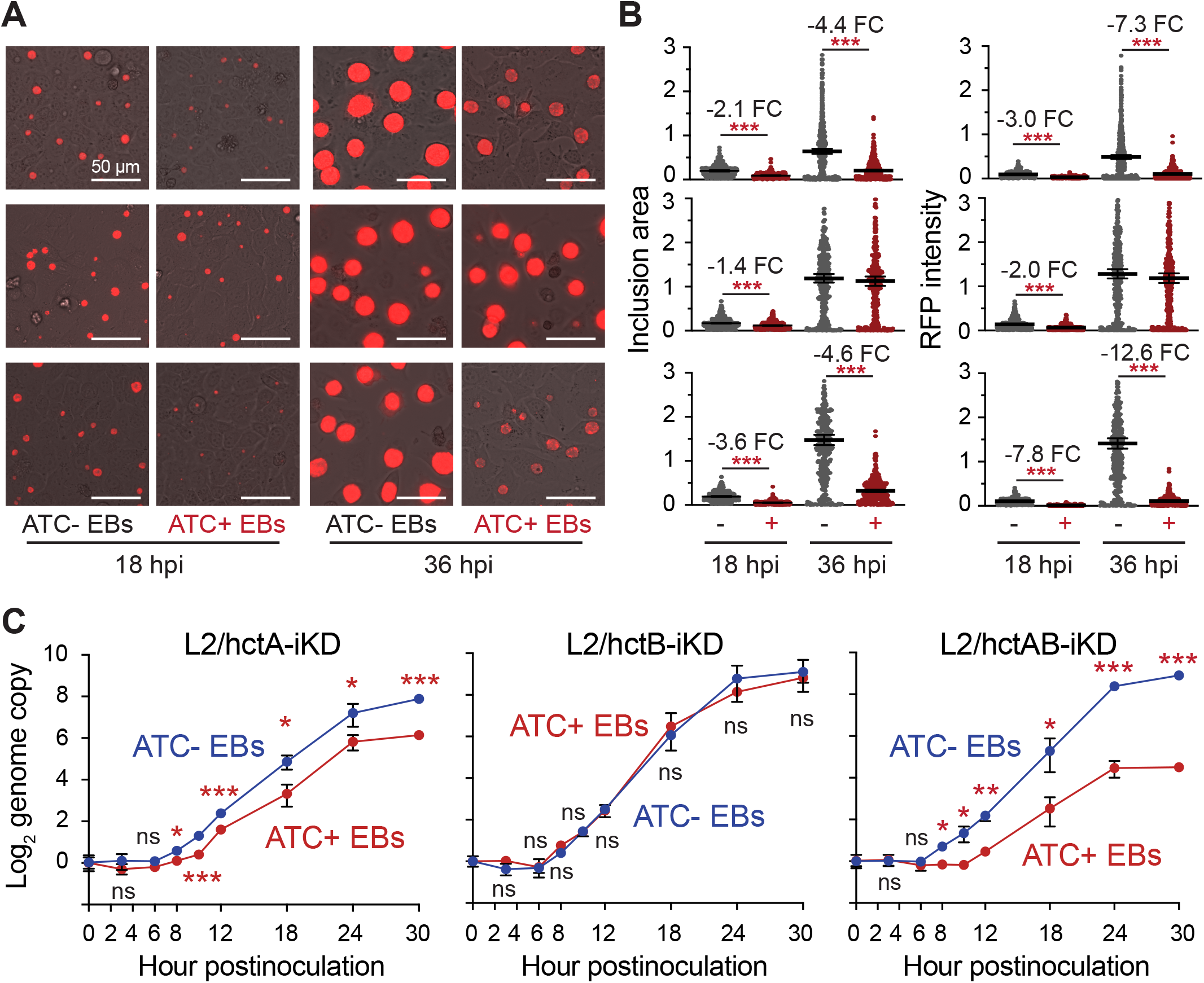
Histone deficiency during EB formation impairs development in secondary infections. (A) Representative fluorescence micrographs of secondary cultures infected with EBs harvested from ATC− or ATC+ primary cultures at 18 and 36 hpi. (B) Quantification of inclusion area and mKate fluorescence intensity in secondary cultures at the indicated time points. “−” and “+” denote EBs harvested from ATC− and ATC+ primary cultures, respectively. (C) Genome copy number over time in secondary cultures infected EBs derived from ATC− or ATC+ primary cultures of L2/hctA-iKD, L2/hctB-iKD, and L2/hctAB-iKD. Data represent averages ± standard deviations from three independent biological replicates. (A-C) EBs were harvested from primary cultures grown with or without ATC and used to infect fresh monolayers in the absence of ATC. Inclusion imaging and genome copy number determination were performed at the indicated times. Data represent three independent biological replicates. Fold changes and statistical significance between conditions are indicated in the graphs.

Quantification of genome copy numbers further resolved these phenotypes mechanistically (Fig. 6C). In cultures inoculated with ATC− EBs, genome copy numbers began to increase by 8 h postinoculation, marking the onset of RB replication. In contrast, cultures inoculated with ATC+ L2/hctA-iKD EBs exhibited little increase through 12 h postinoculation, indicating an approximately four-hour delay in the initiation of detectable replication. After this initial lag, genome accumulation proceeded with reduced kinetics relative to ATC− controls, producing a progressively widening separation between the ATC+ and ATC− trajectories at later time points (Fig. 6C, left). By comparison, genome copy kinetics in cultures inoculated with ATC+ L2/hctB-iKD EBs closely overlapped those of ATC− controls throughout the time course (Fig. 6C, middle). Strikingly, cultures inoculated with ATC+ L2/hctAB-iKD EBs displayed the most severe defects: genome copy numbers remained near baseline until at least 12 h postinoculation and subsequently increased at a substantially reduced rate, resulting in persistently lower genome levels at late time points (Fig. 6C, right). Together, these data indicate that histone deficiency during EB formation imposes a strong transgenerational defect on RB formation and growth, with HctA governing the timing of RB emergence and HctB becoming functionally critical for robust replication when HctA is depleted.

## Discussion

In this study, we used inducible CRISPR interference to define the developmental roles of the chlamydial histones HctA and HctB during late differentiation and across infection cycles. Knockdown of either histone gene, or of both genes simultaneously, produced only modest reductions in EB yield and did not prevent nucleoid condensation during the parental developmental cycle. In striking contrast, EBs generated under histone-deficient conditions, particularly those lacking HctA alone or in combination with HctB, displayed pronounced defects in initiating subsequent infections. These findings indicate that chromatin states established during EB maturation exert durable effects on the next developmental cycle and uncover an unexpected transgenerational dimension to chlamydial chromosome regulation.

The apparent disconnect between relatively mild parental-cycle phenotypes and severe secondary-cycle defects suggests that gross genome condensation and the establishment of developmentally competent chromatin states are not strictly coupled in *Chlamydia*. Even under conditions of substantial histone transcript reduction, EBs formed with condensed nucleoids and retained infectivity. However, those same particles were compromised in their ability to efficiently transition into productive RB growth after reinfection. Genome-replication kinetics in secondary cultures revealed that HctA deficiency primarily delays the onset of RB replication, consistent with impaired EB-to-RB differentiation, whereas combined loss of both histones additionally reduces RB replication capacity at later stages. These observations support a model in which histone-dependent chromatin architecture established during EB maturation governs the efficiency of developmental reprogramming in the next cycle.

Our results further demonstrate that HctA and HctB are not functionally equivalent. Deficiency of HctA exerted a dominant effect on subsequent development, whereas loss of HctB alone produced comparatively mild phenotypes by fluorescence imaging without a corresponding reduction in genome copy number. In contrast, simultaneous knockdown of both histone genes markedly exacerbated growth impairment, indicating that HctB becomes functionally critical when HctA is absent. These distinctions align with prior heterologous-expression studies showing that the two proteins differ in their effects on DNA organization and in their distinct developmental regulation in *Chlamydia*. Together, the data argue for overlapping but non-identical roles, with HctA serving as a principal determinant of chromatin states that prime the genome for reactivation and HctB providing additional structural or regulatory support that becomes essential under conditions of severe chromatin perturbation.

The transgenerational consequences of histone deficiency place HctA and HctB within the broader class of bacterial nucleoid-associated proteins that organize chromosomes and regulate global transcriptomic programs. In diverse bacteria, abundant architectural regulators such as HU, IHF, Fis, and H-NS sculpt large-scale chromosomal domains and couple nucleoid structure to physiological state [21, 22]. Our findings extend this architectural paradigm to a developmental pathogen and demonstrate that chromatin organization established during EB maturation governs fitness in the next infection cycle, manifesting not only as delayed RB reactivation but also as persistently slowed RB replication. Because *Chlamydia* propagates through a differentiation-based developmental program involving morphologically distinct cell types, chromatin-based inheritance may be particularly consequential in this system, ensuring that EBs are not only structurally mature but also primed for efficient progression through the subsequent developmental cycle.

The persistence of nucleoid condensation even after simultaneous knockdown of both histone genes indicates that gross chromatin compaction can still occur under these conditions, although the degree or organization of compaction relative to wild-type EBs remains unknown. This observation suggests that additional factors contribute to EB chromosome packaging or that residual histone protein is sufficient to support visible condensation.

In summary, our results redefine the roles of chlamydial histones beyond structural mediators of EB chromosome condensation. Although HctA and HctB are largely dispensable for EB morphogenesis during the parental developmental cycle, they are essential for establishing chromatin states that support efficient growth in the next round of infection, with HctA playing a dominant role and HctB becoming functionally critical when HctA is depleted. These findings reveal a previously unrecognized layer of epigenetic regulation in *Chlamydia* and establish chromatin organization as a central determinant of developmental fitness across cycles.

## Materials and Methods

### Plasmids

All inducible CRISPRi vectors targeting *hctA, hctB*, or both *hctA* and *hctB* were constructed by assembling PCR fragments amplified from the pL2-idC12-ntg vector [23]. To generate pL2-idC12-hctA, an 8,103-bp fragment was amplified using primers hctA-gRNA-F (5′-ATATGGCGCTAAAAGATACGGCAAATTTTTTTGAAGCTTGGGCC-3′) and mKate-R (5′-AATGCGCATTGTTTGAGTT-3′), and a 7,577-bp fragment was amplified using primers mKate-F (5′-CGTATGAAGGAACTCAAACAAT-3′) and hctA-gRNA-R (5′-ATTTGCCGTATCTTTTAGCGCCATATCTACAAGAGTAGAAATTGAAA-3′). The two fragments were fused using the NEBuilder HiFi DNA Assembly Cloning Kit (New England Biolabs) according to the manufacturer’s instructions.

To generate pL2-idC12-hctB, an 8,103-bp fragment was amplified using primers hctB-gRNA-F (5′-ATATCTATCGACAAGGAGAATGAAATTTTTTTGAAGCTTGGGCC-3′) and mKate-R, and a 7,577-bp fragment was amplified using primers mKate-F and hctB-gRNA-R (5′-ATTTCATTCTCCTTGTCGATAGATATCTACAAGAGTAGAAATTGAAA-3′). The two fragments were assembled using the HiFi DNA Assembly Cloning Kit.

To generate pL2-idC12-hctAB, an 8,187-bp fragment was amplified using primers hctAB-array-F (5′-GGAATTGTGAGCGGATAACAATTTCAATTTCTACTCTTGTAGATATGGCGCTAAAAGATACGGCAAAAATTTCTACTCTTGTAGATATCTATCGACAAGGAGAATGAAA-3′) and mKate-R, and a 7,538-bp fragment was amplified using primers mKate-F and Lac-Operator-R (5′-AATTGAAATTGTTATCCGCTCAC-3′). The fragments were assembled using the HiFi DNA Assembly Cloning Kit.

Complete plasmid sequences of all constructs were confirmed by Nanopore sequencing at Genewiz (Piscataway, NJ).

### Strains

*C. trachomatis* serovar L2 strain 434/BU was obtained from ATCC and propagated in our laboratory in HeLa cells. EBs were transformed with pL2-idC12-hctA, pL2-idC12-hctB, or pL2-idC12-hctAB to generate L2/hctA-iKD, L2/hctB-iKD, and L2/hctAB-iKD, respectively, as previously described [24]. Clonal populations were isolated by limiting dilution [25]; EBs were subsequently purified by ultracentrifugation through 35% and 40%/44%/52% MD-76R density gradients [26].

### *C. trachomatis* infection and culture conditions

HeLa229 cells were maintained in Dulbecco’s modified Eagle medium (DMEM) high-glucose (4.5 g/L) supplemented with 10% fetal bovine serum and 20 µg/mL gentamicin at 37 °C in 5% CO_2_. For gene-expression analyses, growth-rate determinations, and EB-formation assays, cells were seeded into 6- or 12-well plates to form confluent monolayers the following day.

To infect monolayers, EB stocks were diluted in NaHCO_3_-free DMEM/F12 (pH 7.4) to achieve a multiplicity of infection of approximately 1 inclusion-forming unit per cell, yielding 80-90% infection efficiency as assessed by fluorescence microscopy. After removal of the growth medium, EB-containing inoculum was added to the wells (1.5 mL per well for 6-well plates and 1.0 mL per well for 12-well plates). Plates were centrifuged at 900 × g for 30 min at 30 °C and then washed three times with DMEM. Completion of the final wash was defined as 0 hpi.

### RNA isolation and RT-qPCR

Total RNA was extracted from infected cells at 24 hpi using TRI Reagent according to the manufacturer’s instructions. Contaminating genomic DNA was removed by two successive DNase I treatments. One-step RT-qPCR with Luna WarmStart Reverse Transcriptase (New England Biolabs) was used to quantify transcript levels of ddCas12, hctA, hctB, and ihtA. Each sample was analyzed in technical duplicate on a QuantStudio 5 Real-Time PCR System.

Because ATC treatment and histone knockdown did not measurably alter chlamydial genome replication at this time point, 23S rRNA was used for normalization. Differential expression was calculated using the ΔΔCt method [25]. Primer pairs used for ddCas12, hctA, hctB, ihtA, and 23S rRNA were qPCR-dCas12-F (5’-ATGGAACCGTCGTTGAGCTT-3’) and qPCR-dCas12-R (5’-AGGGTCGGCATTTGGAAGTT-3’); qPCR-hctA-F (5’-GGAAATAAAGCCGCAGCAC-3’) and qPCR-hctA-R (5’-ACGATATACCTTCGCGGTCT-3’); qPCR-hctB-F (5’-CATACTGCAGCTTGTGGACG-3’) and qPCR-hctB-R (5’-GCTGTACGAGAACGGTTAGGA-3’); qPCR-ihtA-F (5’-GAGTTGCAAGTTGGTATTCTAACG-3’) and qPCR-ihtA-R (5’-TGTACAAACACTAGAGTCAGAAGC-3’); and qPCR-23S-F (5’-AGATAGACAGCGGGGGCTAA-3’) and qPCR-23S-R (5’-GGTGAGCTGTTACGCACTCT-3’), respectively.

### Sample collection for chlamydial genome copy quantification and IFU assays

At the indicated time points, culture medium was replaced with 500 µL SPG (sucrose– phosphate–glutamic acid solution). Infected cells were detached using cell lifters, collected, and sonicated with a 130-W ultrasonic processor equipped with a 3-mm probe at 35% amplitude for a total of 12 s in alternating 2-s pulses to lyse host cells and release chlamydiae. Lysates were centrifuged at 500 × g to remove host-cell debris. An aliquot (200 µL) of the clarified supernatant was used immediately or stored at −20 °C for genome copy quantification. The remaining suspension was centrifuged at 20,000 × g for 10 min at 4 °C to pellet EBs. Pellets were washed twice with 1 mL SPG to remove residual anhydrotetracycline. After the final wash, pellets were resuspended in 200 µL SPG and stored at −80 °C for subsequent IFU assays.

### Genomic DNA extraction and relative genome quantification

Chlamydial genome copy number was used as a quantitative proxy for total bacterial burden and RB replication. SPG suspensions of chlamydiae were centrifuged at 20,000 × g for 10 min at 4 °C. Pellets were resuspended in 100 µL alkaline lysis buffer (0.1 M NaOH, 0.2 mM EDTA), heated at 95 °C for 15 min, neutralized with 400 µL 25 mM Tris–HCl (pH 7.2), and stored at −20 °C for subsequent analysis, as previously described [23]. Relative chromosome abundance was quantified by qPCR using primers targeting *grgA* (5′-GCCGTTTTTACTGCCAGCAT-3′ and 5′-ACTATTGGAGCCACCTTCGG-3′). Each reaction was performed in technical duplicate on a QuantStudio 5 Real-Time PCR System using Power SYBR Green Master Mix. Genome copy differences were calculated using the ΔCt method.

### IFU assays

IFU assays, analogous to colony-forming unit assays for free-living bacteria, were used to quantify infectious EB production. EB stocks were thawed and tenfold serially diluted in culture medium before inoculation onto 96-well plates containing 3 h-old confluent HeLa229 cell monolayers. Plates were centrifuged at 900 × g for 20 min at 25 °C and then incubated at 37 °C for about 26 h (for plates inoculated with ATC− EBs of all strains and ATC+ EBs of L2/hctB-iKD) or 40 h (for plates inoculated with ATC+ EBs of L2/hctA-iKD and L2/hctB-iKD). Because all chlamydial transformants used in this study express the far-red fluorescent protein mKate, red fluorescent inclusions were enumerated in live cultures using an Olympus IX-50 fluorescence microscope [20].

### Transmission electron microscopy

Ultrastructural analyses were performed essentially as described previously [20]. At 36 hpi, infected HeLa cell monolayers grown in six-well plates were detached using trypsin, collected in phosphate-buffered saline supplemented with 10% fetal bovine serum, and centrifuged for 10 min at 500 × g. Cell pellets were resuspended in fixation buffer containing 2.5% glutaraldehyde and 4% paraformaldehyde in 0.1 M cacodylate buffer at room temperature, incubated for 2 h, and stored at 4 °C overnight. Samples were rinsed in 0.1 M cacodylate buffer, dehydrated through a graded ethanol series, and embedded in Eponate 812 resin at 68 °C overnight. Ninety-nanometer sections were cut using a Leica UC6 ultramicrotome and collected on copper grids. Sections were stained with uranyl acetate followed by lead citrate. Images were acquired as TIFF files on a Philips CM12 transmission electron microscope operated at 80 kV using an AMT XR111 digital camera.

### Fluorescence imaging

EBs harvested from primary cultures grown in the presence or absence of ATC were used to inoculate fresh HeLa monolayers in the absence of inducer. Inclusions were visualized by fluorescence microscopy using the plasmid-encoded mKate reporter present in all strains. Immediately before imaging, culture medium was replaced with phosphate-buffered saline containing calcium and magnesium to reduce background fluorescence. Bright-field and red-fluorescence images were acquired on an Olympus IX51 microscope equipped with an Infinity i8-3 CMOS camera under constant exposure settings. Image overlays were generated using ACINST03 software. The inclusion area and fluorescence intensity were quantified using ImageJ [25].

## ACKNOWLEDGMENTS

This work was supported by National Institutes of Health grants AI140167, AI154305, and AI182210.

